# Omnivory does not preclude strong trophic cascades

**DOI:** 10.1101/441535

**Authors:** Ashkaan K. Fahimipour, David A. Levin, Kurt E. Anderson

**Author notes:** E-mail addresses (Ashkaan K. Fahimipour).

## Abstract

Omnivory has been cited as an explanation for why trophic cascades are weak in many ecosystems, but empirical support for this prediction is equivocal. Compared to predators that feed only on herbivores, top omnivores — species that feed on both herbivores and primary producers — have been observed generating cascades ranging from strong, to moderate, null, and negative. To gain intuition about the sensitivity of cascades to omnivory, we analyzed models describing systems with top omnivores that display either fixed or flexible diets, two foraging strategies that are supported by empirical observations. We identified regions of parameter space wherein omnivores following a fixed foraging strategy, with herbivores and producers comprising a constant proportion of the diet, non-intuitively generate stronger cascades than predators that are otherwise demographically identical: (*i*) high productivity relative to herbivore mortality, and (*ii*) small discrepancies in producer versus herbivore reward create conditions in which cascades are stronger with moderate omnivory. In contrast, flexible omnivores that attempt to optimize *per capita* growth rates during search never induce cascades that are stronger than the case of predators. Although we focus on simple models, the consistency of these general patterns together with prior empirical evidence suggests that omnivores should not be uniformly ruled out as agents of strong trophic cascades.

## Introduction

Trophic cascades occur when top predators indirectly effect change in primary producer biomass by directly reducing populations of intermediate herbivores (Paine, 1980; Strong, 1992; Terborgh & Estes, 2013). A growing number of factors that control the strength of trophic cascades continue to surface from model-based and experimental studies, and their identification has improved our understanding of processes that dampen or enhance indirect effects between species in ecological networks, and ecosystem responses to disturbance (Pace et al., 1999; Shurin et al., 2002; Borer et al., 2005; Shurin et al., 2010; Estes et al., 2011; Heath et al., 2014; Piovia-Scott et al., 2017; Fahimipour et al., 2017). Theories for cascades have traditionally focused on top-down effects in tritrophic food chain models comprising predators that do not directly interact with primary producers (Oksanen et al., 1981; Schmitz et al., 2000; Heath et al., 2014). In many communities however, omnivores that additionally feed on producers occupy top trophic levels (Arim & Marquet, 2004; Thompson et al., 2007). This potential for direct consumption of both producer and herbivore species has led to the prediction, that omnivory should override indirect beneficial effects on producer biomass, thereby dampening or disrupting cascades in most cases (Polis & Strong, 1996; Pace et al., 1999; Bruno & O’Connor, 2005; Duffy et al., 2007; Kratina et al., 2012; Wootton, 2017).

Documented instances of weakened and even reversed trophic cascades in food webs with omnivory (Flecker, 1996; Pringle & Hamazaki, 1998; Snyder & Wise, 2001; Finke & Denno, 2005; Bruno & O’Connor, 2005; Denno & Finke, 2006; Johnson et al., 2014; Visakorpi et al., 2015; Fahimipour & Anderson, 2015), compared to those typically induced by predators (Shurin et al., 2002), are not uncommon and provide support to the intuitive hypothesis that omnivory precludes strong trophic cascades. However, a large meta-analysis of 114 experimental predator and omnivore manipulations in terrestrial, freshwater, and marine systems could not identify differences in the magnitudes of trophic cascades between the two groups (Borer et al., 2005). Empirical evidence to the contrary — namely, examples of stronger or comparable cascades that are generated by omnivores (Power, 1990; Power et al., 1992; Okun et al., 2008; France, 2012) — implies that weak cascades may not be a guaranteed outcome of omnivory in food webs. Despite a growing body of theoretical and empirical work, an understanding of when omnivores occupying top trophic positions will generate strong or weak cascading effects is lacking, and likely depends on multiple population- or community-level factors (Wootton, 2017).

We analyzed mathematical models describing trophic interactions between basal producers, intermediate herbivores, and top omnivores to systematically evaluate the effects of omnivory on the strength of trophic cascades. We consider two types of empirically-observed foraging behaviors, namely *fixed* (Diehl & Feißel, 2000) and *flexible* (Fahimipour & Anderson, 2015) omnivory (see *Model Formulations* for definitions), and present a comparison between trophic cascades in these systems and traditional ones induced by analogous predators. We have chosen to study minimally detailed models to focus on coarse-grained system features that may point to potential future directions for experimental work, as opposed to making predictions about the behavior of a particular ecosystem (Anderson et al., 2009). We draw two primary conclusions based on numerical and analytical results: stronger trophic cascades with omnivory are at least possible in high productivity systems if omnivores forage according to a fixed strategy, whereas cascades are never stronger when omnivores forage according to a flexible strategy.

## Model Formulations

Models were analyzed with a focus on equilibrium outcomes to gain insight into how differences in the foraging strategies of species occupying top trophic levels (i.e., predators versus fixed or adaptive omnivores) influence long-term community structure as measured by the trophic cascade. We modeled the population dynamics of three species: (*i*) basal producers, that are eaten by (*ii*) intermediate herbivores and (*iii*) top omnivores that consume both producers and herbivores (Diehl & Feißel, 2000). Analyses of similar three-node trophic modules have demonstrated how the coexistence of all species and community stability are sensitive to variation in system primary productivity and the strength of omnivory (parameters *ρ* and *ω* in eqs. 1 and 3 below; discussed extensively by McCann & Hastings, 1997; Diehl & Feißel, 2000, 2001; Gellner & McCann, 2011). For this reason, a primary goal of our analysis was to elucidate how primary productivity and omnivory strength interact to influence trophic cascades in three species modules with and without true omnivory.

Two omnivore foraging strategies with empirical support were considered. We refer to the first as a *fixed* foraging strategy, indicating that foraging effort toward either producers or herbivores comprise constant proportions of the *fixed omnivores*’ total foraging effort (McCann & Hastings, 1997; Diehl & Feißel, 2000). The second strategy, which we refer to as *flexible* foraging, indicates that the foraging effort apportioned toward either producers or herbivores by the *flexible omnivore* varies in time, and depends on the availability and reward associated with each resource species (Kondoh, 2003).

### Fixed foragers

We assume a linear (type I; Holling, 1959) functional response relating resource densities to *per capita* consumption rates, so that the dynamics of species’ biomasses are represented by the system of equations

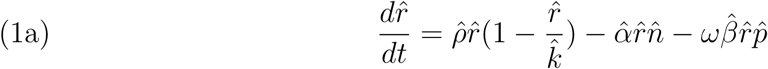

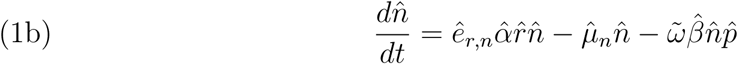

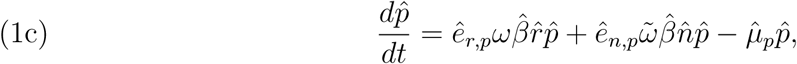

where hats over terms indicate that they have dimensions, and 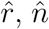, and 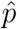 are the biomasses of producers, herbivores, and omnivores.

Here, 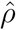 and 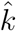 are the producer productivity rate and carrying capacity, 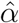 is the herbivore foraging rate, 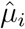 is the *per capita* mortality rate of species *i*, and *ê*_*i,j*_ is the resource *i* assimilation efficiency for consumer *j*. We assumed a total foraging rate 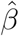 for omnivores, that is apportioned toward herbivores proportionately to 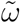, where 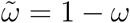. We therefore interpret *ω* as a nondimensional parameter describing omnivory strength (McCann & Hastings, 1997); the system reduces to a food chain when *ω* = 0. See Table 1 for a summary of all model parameters.

**Table 1.** Parameter descriptions for equations (1) and (2).

| Parameter | Description | Units | Range or Value |
| --- | --- | --- | --- |
| $\hat{\rho}$ | producer productivity rate | time<sup>-1</sup> | $\hat{\rho} > 0$ |
| $\hat{k}$ | producer carrying capacity | producer · area<sup>-1</sup> | $\hat{k} > 0$ |
| $\hat{\alpha}$ | herbivore foraging rate | area · herbivore<sup>-1</sup> · time<sup>-1</sup> | $\hat{\alpha} > 0$ |
| $\hat{\beta}$ | omnivore foraging rate | area · omnivore<sup>-1</sup> · time<sup>-1</sup> | $\hat{\beta} > 0$ |
| $\omega$ | fraction of omnivore foraging effort toward producers | dimensionless | $0 < \omega < 1$ |
| $\tilde{\omega}$ | fraction of omnivore foraging effort toward herbivores | dimensionless | $1 - \omega$ |
| $\hat{e}_{i,j}$ | conversion efficiency of resource $i$ to consumer $j$ | units of $j$ · units of $i$<sup>-1</sup> | $0 < \hat{e}_{ij} < 1$ |
| $\hat{\mu}_i$ | mortality rate of species $i$ | time<sup>-1</sup> | $\hat{\mu}_i > 0$ |
| $v$ | time scale of behavioral change | dimensionless | $v > 0$ |

### Flexible foragers

Equations (1) can be modified to include flexible foraging behavior by the omnivore, by substituting the omnivory strength parameter *ω* with the dynamical state variable Ω. Flexible foraging behavior was modeled using a replicator-like equation (Kondoh, 2003), which provides a reasonable representation of flexible omnivory in real food webs (Fahimipour & Anderson, 2015). The behavioral equation is

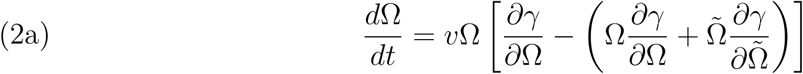

where 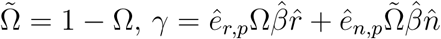 is the flexible omnivore’s instantaneous *per capita* biomass production rate, and the constant *v* is a nondimensional ratio between the time scales of foraging adaptation and omnivore population dynamics (Heckmann et al., 2012). Values of *v* > 1 represent behavioral changes that occur on faster time scales than omnivore generations. This behavioral model implies that omnivores gradually adjust their foraging strategy during search if behavioral changes yield a higher instantaneous *per capita* biomass production rate than the current diet (Kondoh, 2003).

### Model nondimensionalizations and assumptions

The parameters in equations (1) and (2) were transformed into nondimensional parameters using scaled quantities, reducing the total number of model parameters to those with values having clear interpretations (Murray, 1993; Nisbet & Gurney, 2003). We use substitutions similar to Amarasekare (2006) and Amarasekare (2007):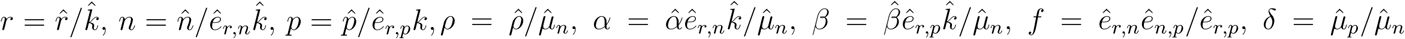, and 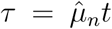. Substituting into eqs. (1) and (2), we obtain the nondimensional system

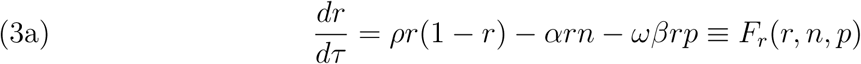

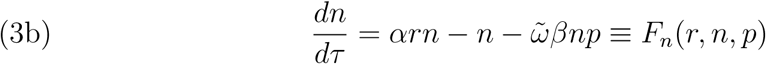

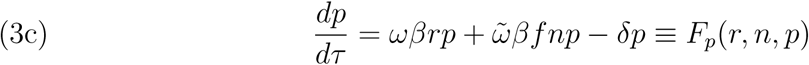

for fixed omnivory, with

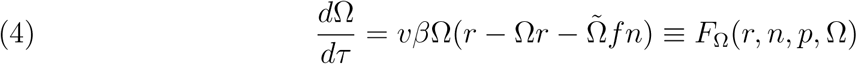

representing flexible foraging behavior. For the fixed model, scaled producer, herbivore, and omnivore biomasses are represented as ***x*** = [*r, n, p*]. The vector field which maps [*r, n, p*] to [*F*_*r*_(*r, n, p*), *F*_*n*_(*r, n, p*), *F*_*p*_(*r, n, p*)] is denoted by *F*_fixed_ : ℝ ^3^ →ℝ ^3^, and the coexistence equilibrium of the fixed foraging model eq. (3) is denoted by 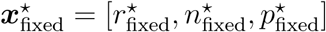. We considered trophic cascades in systems with stable equilibria, satisfying

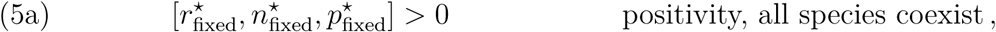

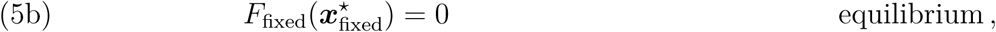

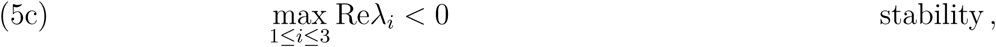

where 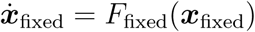 describes the system of equations (3), and *λ*_*i*_ are the eigenvalues of the Jacobian matrix evaluated at equilibrium 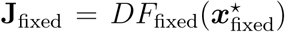. These conditions ensure a straightforward comparison of trophic cascades, which in the case of nonstationary steady states would depend on the time scales under consideration (Borer et al., 2005).

Eq. (3) is extended to the case of flexible foraging by replacing the fixed foraging parameter *ω* with a quantity satisfying (4). The system of equations is now four-dimensional and is defined by the vector field *F*_flexible_ : ℝ^4^ → ℝ^4^ which maps [*r, n, p*, Ω] to [*F*_*r*_(*r, n, p*, Ω), *F*_*n*_(*r, n, p*, Ω), *F*_*p*_(*r, n, p*, Ω), *F*_Ω_(*r, n, p*, Ω)]. We demand that the flexible model likewise has a coexistence equilibrium 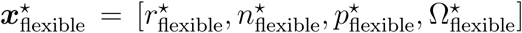, so that 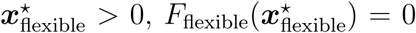, and all eigenvalues of the system’s Jacobian matrix 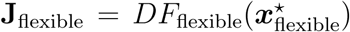 have negative real parts. Finally, for the case of predators in a food chain, that do not feed on primary producers, we denote by 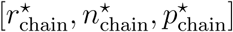 the stable and positive solution satisfying (5) when setting *ω* = 0.

### A comparison of trophic cascades

We quantified differences in trophic cascade strengths between systems with omnivores (i.e., *ω* > 0) and predators (i.e., *ω* = 0), and examined the dependencies of these differences on model parameters. We denote by 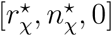 the non-positive equilibrium solution to (3) in the absence of predators, so that 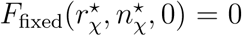. A traditional measure of trophic cascade strength (Shurin et al., 2002; Borer et al., 2005) applied to omnivory systems at equilibrium is therefore 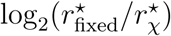. Likewise, cascade strength in the analogous food chain can be calculated as 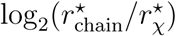. The difference in trophic cascade strengths induced by a fixed omnivore and the predator in its analogous food chain, *κ*_*fixed*_

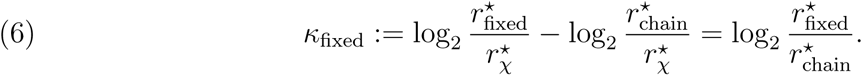

This measure *κ*_*fixed*_ of the relative cascade strength is similar to the “proportional response” measure of Heath et al. (2014), and equals 1 (or *-*1) if the trophic cascade induced by omnivores is twice as strong (or half the strength) as in the analogous food chain. Like-wise, the difference in cascade strengths between flexible omnivory systems and food chains, 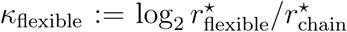. Closed form equilibrium solutions for all variables in eqs. (3) and (4) are provided in Supplementary Table 1.

## Results

### Fixed omnivores can generate strong trophic cascades

We first considered the case of fixed omnivores that do not exhibit diet flexibility. In Figure 1 we summarize changes in the relative cascade strengths induced by fixed omnivores *κ*_fixed_ (Fig. 1a), community dynamics, and species coexistence (Fig. 1b) as primary productivity *ρ* and omnivory strength *ω* — two key determinants in the behavior of omnivory systems (McCann & Hastings, 1997; Diehl & Feißel, 2001; Amarasekare, 2007; Gellner & McCann, 2011) — are varied while other parameters are held constant. Consistent with prior analyses of three-species omnivory modules, increasing omnivory strength *ω* causes the system to undergo a transcritical bifurcation (McCann & Hastings, 1997) resulting in extinction of either omnivores at low productivities, or herbivores at high productivities (Fig. 1b; Diehl & Feißel, 2001; Amarasekare, 2007). These occur when the determinant of the Jacobian matrix vanishes, det **J**_fixed_ = 0, for a combination of *ρ* and *ω*, marking the presence of a zero eigenvalue (Kuznetsov, 2013).

**Figure 1.**
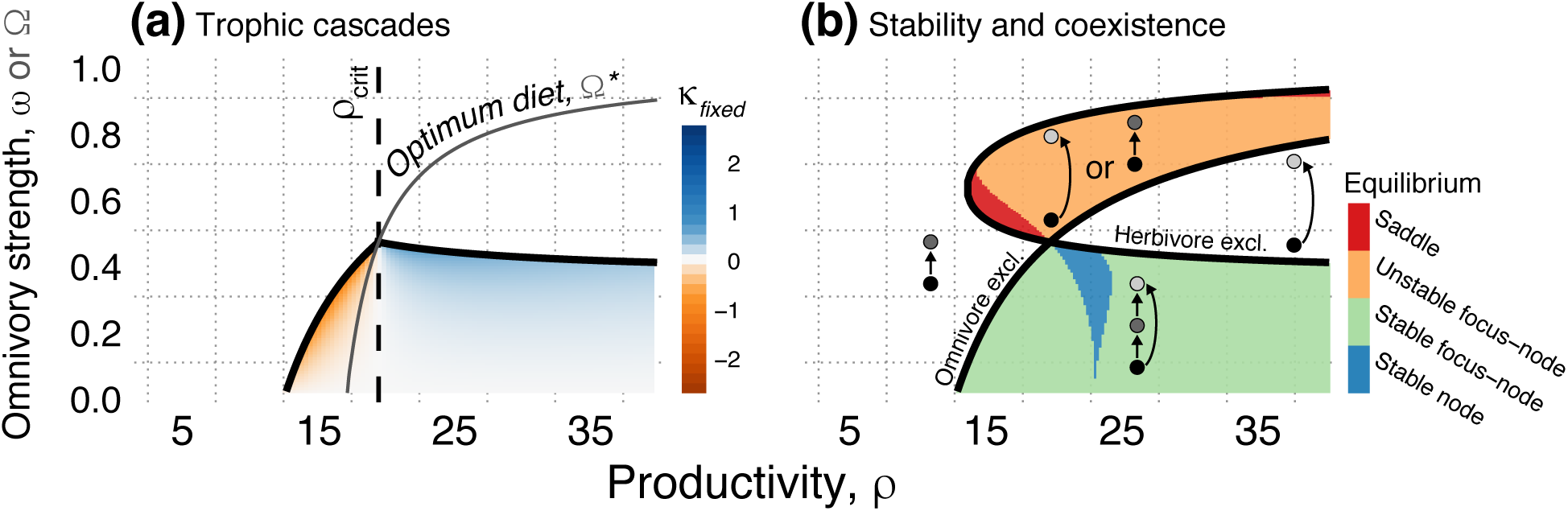
(a) Numerical summary of relationships between relative cascade strength *κ*_*fixed*_, productivity *ρ*, and omnivory strength *ω*. Colors show *κ*_*fixed*_ values for combinations of *ρ* and *ω* within the three-species coexistence region. Blue represents stronger cascades with omnivory, orange represents weaker cascades with omnivory. Black curves mark extinction boundaries for either the omnivore or herbivore species (see panel b). The grey curve shows the equilibrium foraging strategy for the flexible omnivore, 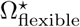. A vertical dashed line marks the critical productivity *ρ*_crit_. (b) Bifurcation curves identify stability boundaries separating steady states with different dynamics. Colors are different types of equilibria, determined by eigenvalues of the Jacobian matrix **J**_fixed_. Non-coexistence regions are labelled and identified by the inset networks of producers (black), herbivores (dark grey), and omnivores (light gray). Parameter values are *α* = 7.5, *β* = 5.5, *f* = 0.25, *δ* = 2, and *v* = 1.05.

Within the stable coexistence region (Fig. 1b, green and blue regions), predictions of weaker trophic cascades with omnivory (Pace et al., 1999; Shurin et al., 2010; Kratina et al., 2012; Wootton, 2017) held when primary productivity, *ρ*, was below a threshold value (Fig. 1a, orange region). Productivities above this threshold however, yield omnivory cascades that are non-intuitively stronger compared to predators (Fig. 1a, blue region). This critical transition in relative trophic cascade strengths with increasing productivity occurs at a point which we refer to as *ρ*_crit_ or the *critical productivity* for convenience (Fig. 1a). A vertical dashed line marks the critical productivity, which is the value of *ρ* at which *κ*_fixed_ = 0, given parametrically by *ρ*_crit_ = *δα*^2^(*f -* 1)*/f* [*α*(*δ - β*) + *βf* (*α -* 1)] (Supplementary Fig. 1). Note that the transition from weaker (*κ*_*fixed*_ < 0) to stronger (*κ*_*fixed*_ > 0) cascades along a productivity gradient does not depend on omnivory strength. Instead, omnivory strengths near the extinction boundaries attenuate the discrepency between cascades, such that omnivory cascades are weakest when productivity is low and *ω* approaches values leading to omnivore exclusion, and strongest when productivity is high and omnivores have nearly excluded herbivores (Fig. 1).

To explain the non-intuitive result of stronger cascades with fixed omnivory, we examined the relationship between primary productivity and the optimal foraging effort that would lead to the highest *per capita* growth rate by omnivores at equilibrium, 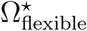 (Supplementary Table 1). The grey curve in Fig. 1a illustrates 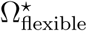 as a function of *ρ*; the growth rate-maximizing strategy monotonically approaches pure herbivory as productivity increases, recapitulating results that coexistence occurs over a wider range of omnivory strengths when omnivores forage flexibly (Křivan & Diehl, 2005). Precisely at *ρ* > *ρ*_crit_, the fixed omnivore is no longer able to achieve the optimal foraging strategy (Fig. 1a). Intuitively, this indicates that strong trophic cascades are induced by omnivores when their foraging effort toward producers is guaranteed to be energetically suboptimal.

We next sought to determine whether the presence of a critical productivity, or a switch from weaker to stronger cascades with fixed omnivory, depends on other model parameters. In Fig. 2 we show that *ρ*_crit_ (i.e., the location of the vertical dashed line in Figure 1a along the *x*-axis) is sensitive to the conversion efficiency parameter, *f*. Recall that *f* is the product of the producer-to-herbivore conversion efficiency ê_*r,n*_, and the ratio of herbivore- and producer-to-omnivore conversion efficiencies (i.e., omnivore rewards), ê_*n,p*_*/*ê_*r,p*_ (Table 1). Model-based and experimental studies have suggested that ê_*r,n*_ > ê_*r,p*_ and ê_*n,p*_ > ê_*r,p*_ are realistic conditions for omnivores in nature (Diehl & Feißel, 2000; Křivan & Diehl, 2005). Satisfying these conditions, we would generally expect small *f* values when herbivores are only slightly more rewarding than producers to omnivores (for instance, if ê_*n,p*_ = ê_*r,p*_ + *ϵ* where ϵ is a small number), and large *f* values when rewards from eating herbivores are much higher than for producers, ê_*n,p*_ ≫ ê_*r,p*_. For large enough values of *f*, the curve of *ρ*_crit_ enters a non-coexistence region (Fig. 2a). Thus, the potential for strong omnivory cascades is lost as *f* increases, regardless of other population- or community-level properties; an example of when this happens is shown in Figs. 2c and 2e. We examine the sensitivity of these results to other model parameters in Supplementary Fig. 1. Briefly, the critical productivity shifts to the right (i.e., larger *ρ*_crit_ values lead to smaller parameter regions with strong omnivory cascades) as *α* and *δ* increase, and shifts toward zero (i.e., smaller *ρ*_crit_ values lead to an expansion of the parameter region with stronger omnivory cascades) as *β* increases (Supplementary Fig. 1).

**Figure 2.**
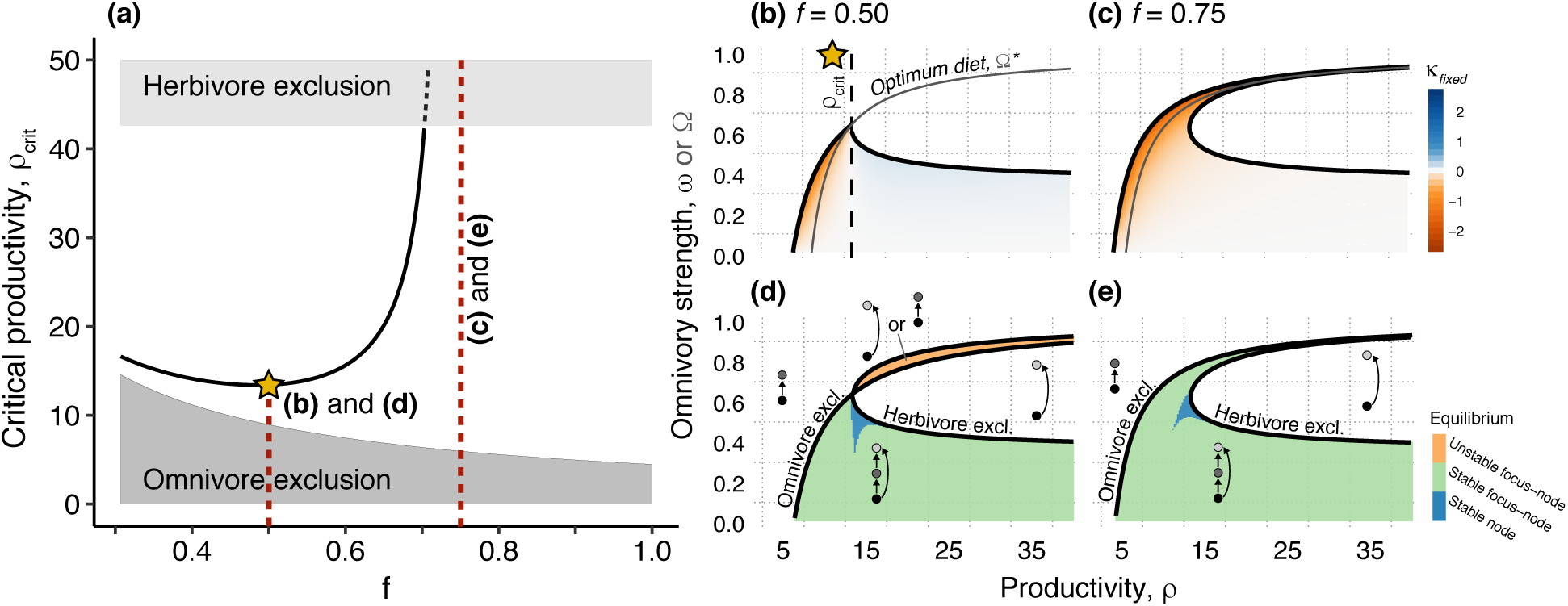
(a) Relationship between critical productivity *ρ*_crit_ and efficiencies *f* (Table 1). The curve is solid if the critical productivity lies in the coexistence region, and dashed otherwise. The light and dark grey shaded regions mark the extinction of herbivores and omnivores respectively. The left- and righthand red dashed lines correspond to panels b & d and c & e respectively. Parameter values are the same as in Fig. 1. (b & c) Relationships between relative cascade strength *κ*_*fixed*_, productivity *ρ*, and omnivory strength *ω*. Colors show *κ*_*fixed*_ values within the three-species coexistence region. Blue represents stronger cascades with omnivory, orange represents weaker cascades with omnivory. Black curves mark extinction boundaries for either the omnivore or herbivore species (see panels d & e). The grey curve shows the equilibrium foraging strategy for the flexible omnivore, 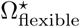. A vertical dashed line marks the critical productivity *ρ*_crit_. (d & e) Bifurcation curves identify stability boundaries separating steady states with different dynamics. Colors mark different types of equilibria. Non-coexistence regions are labelled and identified by the inset networks of producers (black), herbivores (dark grey), and omivores (light gray).

### Flexible omnivores never generate stronger trophic cascades

Unlike fixed omnivores, flexibly foraging omnivores can never induce cascades that are stronger than in the analogous food chain. We show analytically that at a positive equilibrium solution, *κ*_*F*_ < 0. At the interior equilibrium (Supplementary Table 1), if *ϕ* := –*αδ* + *δf* (*α* – *ρ*) + *βf* (*ρ* – 1) then

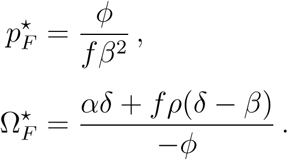

The ratio of flexible omnivory to linear chain trophic cascade strengths,

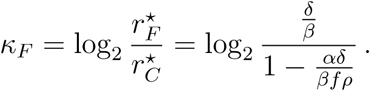

As 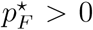 and evidently *fβ*^2^ > 0, we must have *ϕ* > 0. Moreover, since 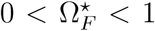, we must also have *δfρ* > *βfρ* – *αδ*. That is,

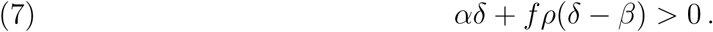

Combining (7) with the solution 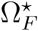 (Supplementary Table 1) shows that for positive equilibria, *κ*_*F*_ < 0, since *ϕ* < 0 cannot be true for a biological system. Thus, consistent with conceptual models of trophic cascades (Strong, 1992; Pace et al., 1999), cascades in systems with flexibly foraging top omnivores are bounded in strength by those in their analogous food chains. Numerical results confirm these analytical expectations, and illustrate how in-creasing consumer reward ratios (i.e., increasing *f*) attenuates this result but does not alter the qualitative relationship between *κ*_flexible_ and *ρ* (Fig. 3).

**Figure 3.**
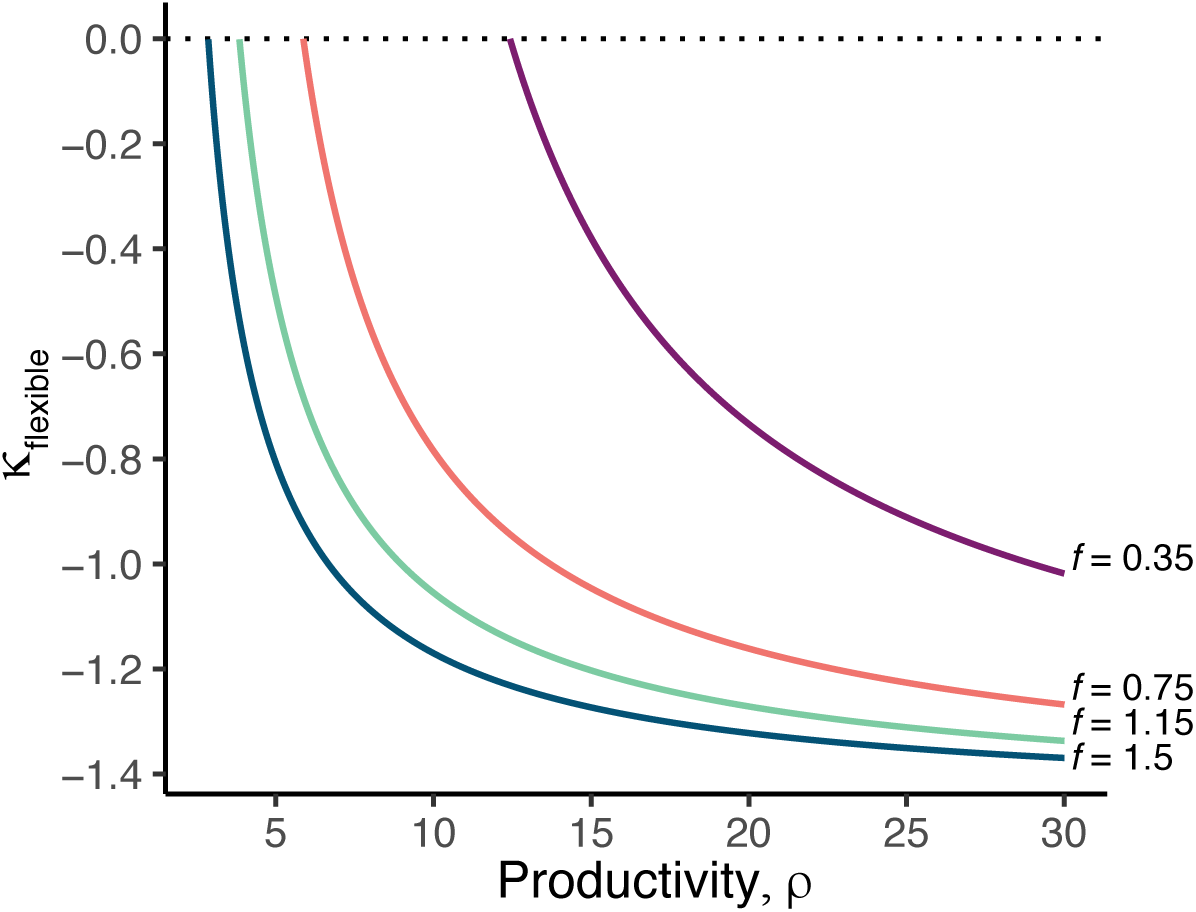
Numerical solutions relating relative cascade strength, *κ*_*F*_, scaled productivity *ρ*, and scaled resource profitability *f* in the flexible model. Colors represent associated values of *f* in the margin. Parameter values are *α* = 7.5, *β* = 5.5, *δ* = 2, and *v* = 1.05.

## Discussion

Intuition suggests that trophic cascade will not occur when top predators additionally feed on primary producers (Polis & Strong, 1996; Pace et al., 1999; Duffy et al., 2007; Shurin et al., 2010; Kratina et al., 2012; Wootton, 2017), but our results predict that strong cascades will emerge under a wider range of foraging types than previously expected. We identified many cases in which omnivores are indeed likely to generate weak cascades, although we have shown that this should not be a uniform expectation for omnivory. Particularly, in high productivity systems in which forging rewards do not strongly differ between producers and herbivores (Fig. 1a; Fig. 2a), fixed omnivores are capable of generating stronger cascades than would be expected if they did not consume producers at all. This is due to suboptimal omnivore foraging, and the additional source of herbivore population losses in models of fixed omnivory, in which the herbivore must compete with its own consumer for resources (Diehl & Feißel, 2000). This result provides at least one general explanation for the weaker (Finke & Denno, 2005; Denno & Finke, 2006), comparable or indistinguishable (Borer et al., 2005), and stronger (Okun et al., 2008; France, 2012) cascades that have now been observed with omnivorous top predators: they largely depend on primary productivity and the types of omnivory. It is not surprising that a more comprehensive catalogue of foraging behaviors will improve predictions of trophic cascades, but our model-based results indicate that this knowledge may be especially important when species consume resources across trophic levels.

Comparisons of fixed and flexible models showed that omnivores were capable of generating strong cascades only when consuming an energetically suboptimal level of primary producers could be guaranteed (Figs. 1a, 2b, and 2c). This leads to the question: how common is this type of fixed foraging in food webs? Empirical evidence for approximately fixed foraging exists for groups as diverse as protists, arthropods, and mammals (Clark, 1982; Mooney & Tillberg, 2005; Diehl & Feißel, 2001). Fixed omnivory may also manifest in other ways, for example when organisms forage in a way that is suboptimal in terms of pure energetics but is otherwise required to maintain nutritional or stoichiometric balances (Berthoud & Seeley, 1999; Remonti et al., 2016; Zhang et al., 2018). Suboptimal foraging has also been observed in heavily disturbed or human-altered systems where consumer behaviors are not adapted to current resource conditions, or when changes in habitat structure alter the ability to efficiently locate preferred food sources (Walsh et al., 2006).

Allometric scaling relationships between species’ demographic rates and body masses have helped identify biological constraints on the strengths of trophic cascades in food chains with top predators (DeLong et al., 2015), but body mass may have additional implications for cascades that are generated by species facing complex foraging decisions. The prevalence of dynamical or adaptive foraging behaviors, like those represented by our flexible model, across the tree of life has shown associations with organismal brain sizes and body masses by proxy (Eisenberg & Wilson, 1978; Rooney et al., 2008; Edmunds et al., 2016). Body mass distributions may also influence cascades that are induced by species with size-mediated ontogenetic shifts from herbivory to carnivory (Pace et al., 1999), wherein average population-level foraging behaviors could be characterized as “omnivory” and would reflect intraspecific size structures. Future empirical work and simulation-based analyses of more complex models will be key for uncovering additional relationships between species’ body masses and trophic cascades in food webs, and to develop a coherent understanding of when foraging behavior drives deviations from predictions of cascades from simple tritrophic food chain models. In many of these case, omnivory could appear as an average population-level behavior and not necessarily at the level of the individual.

Our analysis focuses on models characterized by type I functional responses that relate resource biomass to consumer growth. Alternate nonlinear functional responses (e.g., Holling type II; Holling, 1959) may modulate the effects of omnivory on trophic cascades. Preliminary analyses show that closed-form equilibrium solutions similar to those in Supplementary Table 1 can also be obtained for Type II functional responses. The predictive power of these solutions in cases where the model shows oscillatory behavior remains an open question. We conjecture that, for mild instabilities, the oscillatory behavior introduced by saturating consumption would result in similar qualitative outcomes predicted by equilibrium values when cascades are measured as time-averaged quantities (Fox, 2007). However, for larger-amplitude oscillations cascade strengths will likely depend strongly on the time-scale over which they are measured, potentially yielding a mechanism for the observation that cascade strengths are related to experimental duration (Borer et al., 2005). Models that formally extend the concept of trophic cascades to cases of nonstationary equilibria will be an important direction for future analyses.

Figure 1 suggests an interesting analytical question for future study. Namely, multiple qualitative changes are observed precisely at the phase transition for strong omnivory cascades, *κ*_fixed_ > 0, which is indicated by the vertical dashed line *ρ* = *ρ*_crit_. Also occurring at this point are the phase boundaries for species coexistence at stable equilibrium given by the curves separating different non-coexistence regions (Fig. 1b); the grey curve showing the optimal omnivory strength, 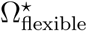, as a function of *ρ* passes into a region that is unattainable by the fixed forager (Fig. 1a); and two saddle-node bifurcation curves intersect (Fig. 1b). It remains to understand why these curves all intersect at a single point, and how this relates to qualitative changes in trophic cascade strengths.

Examples from agroecosystems and disturbed natural habitats indicate that trophic cascade theories can directly inform applied management problems and efforts to mitigate human alteration of ecosystems (Schmitz, 2006; Estes et al., 2011). Our comparative analyses together with the ubiquity of omnivory in nature (Arim & Marquet, 2004; Kratina et al., 2012; Wootton, 2017) suggest that omnivores may contain promise for such applications of cascade theory. For instance, nutrient inputs to agricultural systems that lead to artificially enriched communities are exactly the conditions where we expect a potential for strong omnivorous cascades. If management goals include reducing the density of agricultural pests in enriched systems through integrated strategies that manipulate top trophic levels, then, counterintuitively, top omnivores with certain features may warrant additional consideration (Agrawal et al., 1999). Achieving these outcomes in practice may prove challenging (Cortez & Abrams, 2016).

### Conclusions

Omnivory has long been cited as a reason for why trophic cascades are less frequent or weaker than expected, although empirical data on the role of omnivory has been equivocal (Borer et al., 2005; Shurin et al., 2010; Kratina et al., 2012; Wootton, 2017). Our theory generally agrees with the prediction of omnivory in weakening cascades, but also demonstrates where these predictions are weak or even where they exhibit unexpected changes. Thus, these predictions generate a framework for future investigation, that can focus expectations on when and where omnivory effects might occur in more complex ecosystems. At the least, our models help elucidate the mixed support for an intuitive ecological prediction.

## Acknowledgements

We thank two reviewers for comments. A.K.F was supported by a grant from the Alfred P. Sloan Foundation to the Biology & the Built Environment Center at the University of Oregon, and by the National Science Foundation (NSF) award #006741-002. K.E.A. was supported by NSF grant DEB #1553718.

## Supplementary Materials

**Supplementry Table 1.**
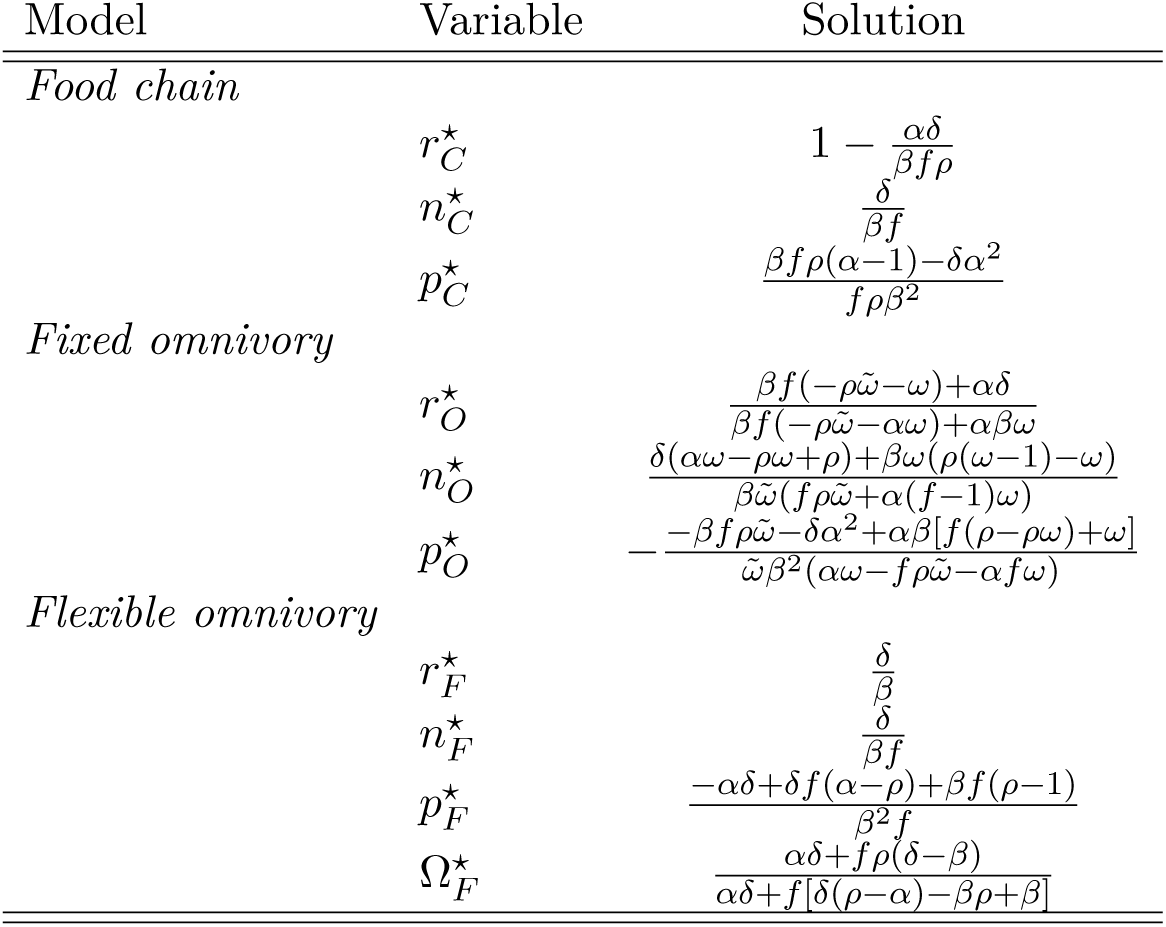
Closed-form equilibrium solutions for all systems of equations. Variables and parameters in the *Variable* and *Solution* columns refer to the nondimensional scaled quantities in eqs. (3) and (4).

| Model | Variable | Solution |
| --- | --- | --- |
| *Food chain* | $r_C^*$ | $1 - \frac{\alpha\delta}{\beta f \rho}$ |
| | $n_C^*$ | $\frac{\delta}{\beta f}$ |
| | $p_C^*$ | $\frac{\beta f \rho(\alpha-1) - \delta \alpha^2}{f \rho \beta^2}$ |
| *Fixed omnivory* | $r_O^*$ | $\frac{\beta f(-\rho\tilde{\omega}-\omega) + \alpha\delta}{\beta f(-\rho\tilde{\omega}-\alpha\omega) + \alpha\beta\omega}$ |
| | $n_O^*$ | $\frac{\delta(\alpha\omega - \rho\omega + \rho) + \beta\omega(\rho(\omega-1) - \omega)}{\beta\tilde{\omega}(f\rho\tilde{\omega} + \alpha(f-1)\omega)}$ |
| | $p_O^*$ | $-\frac{-\beta f \rho\tilde{\omega} - \delta\alpha^2 + \alpha\beta[f(\rho - \rho\omega) + \omega]}{\tilde{\omega}\beta^2(\alpha\omega - f\rho\tilde{\omega} - \alpha f\omega)}$ |
| *Flexible omnivory* | $r_F^*$ | $\frac{\delta}{\beta}$ |
| | $n_F^*$ | $\frac{\delta}{\beta f}$ |
| | $p_F^*$ | $\frac{-\alpha\delta + \delta f(\alpha - \rho) + \beta f(\rho - 1)}{\beta^2 f}$ |
| | $\Omega_F^*$ | $\frac{\alpha\delta + f\rho(\delta - \beta)}{\alpha\delta + f[\delta(\rho - \alpha) - \beta\rho + \beta]}$ |

**Supplementry Figure 1.**
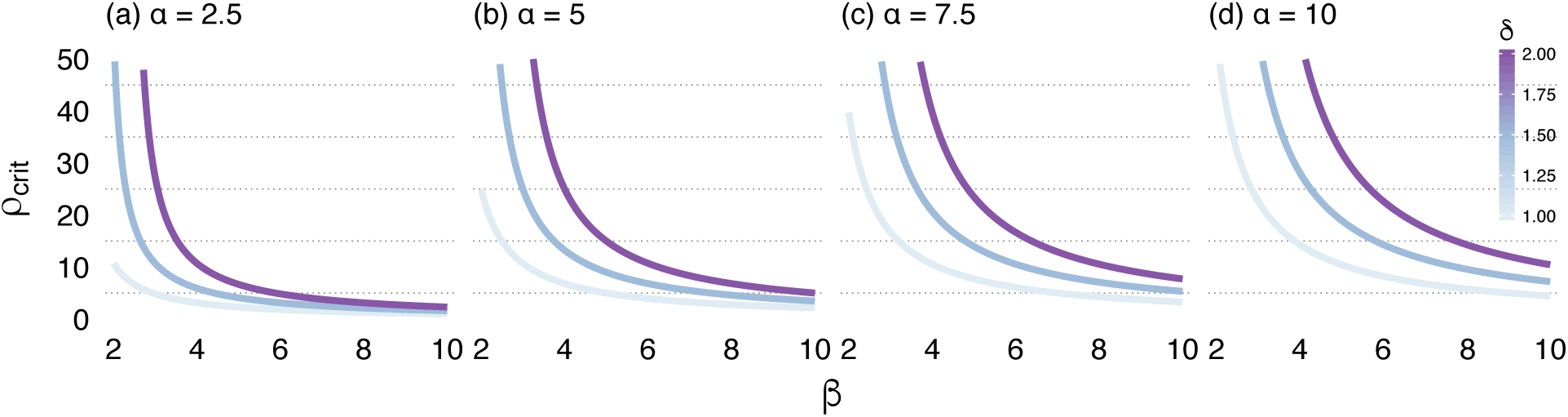
The critical productivity *ρ*_crit_ as a function of *α* (panels), *β* (*x*-axis), and *δ* (color scale). The parameter *f* = 0.5.

