## Supplementary figures and images for "Omnivory does not preclude strong trophic cascades"

### Supplemental Figure 1

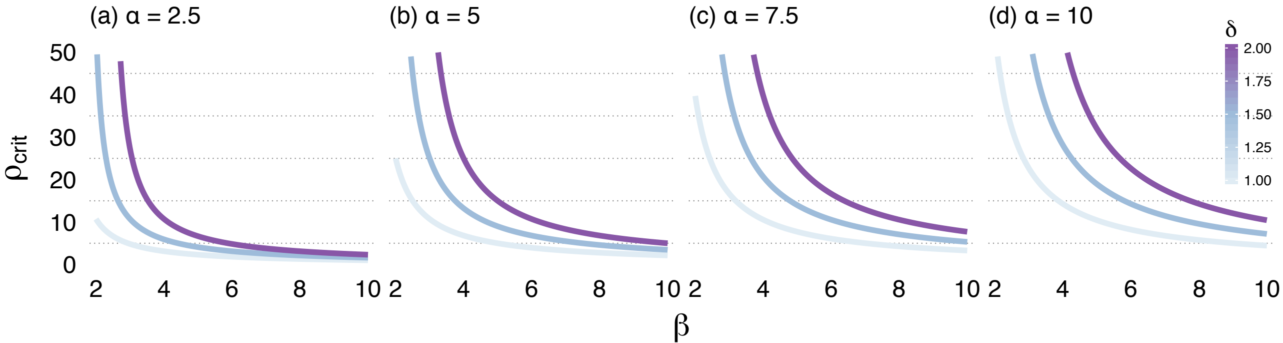
